# Extensive MET alterations confer clinical response to MET inhibitors in gliomas

**DOI:** 10.1101/2020.11.02.364711

**Authors:** Zheng Zhao, Jing Chen, Zhaoshi Bao, Ruichao Chai, Ke-nan Zhang, Lingxiang Wu, Hanjie Liu, Quanhua Mu, Huimin Hu, Fan Zeng, Zheng Wang, Guanzhang Li, Yuanhao Chang, Qiangwei Wang, Fan Wu, Ying Zhang, Yuqing Liu, Wei Zhang, Chunsheng Kang, Jiguang Wang, Rongjie Tao, Qianghu Wang, Tao Jiang

**Affiliations:** Beijing Neurosurgical Institute, Capital Medical University, Beijing, China; Department of Neurosurgery, Beijing Tiantan Hospital, Capital Medical University, Beijing, China; Department of Bioinformatics, Nanjing Medical University, Nanjing, Jiangsu, China; Departments of Neurosurgery, Shandong Cancer Hospital and Institute, Shandong University, Jinan, Shandong, China; Department of Chemical and Biological Engineering, The Hong Kong University of Science and Technology, Hong Kong, China; Department of Neurosurgery, Tianjin Medical University General Hospital, Tianjin, China; Center of Brain Tumor, Beijing Institute for Brain Disorders, Beijing, China; China National Clinical Research Center for Neurological Diseases, Beijing, China

## Abstract

Activating alterations of the MET gene are well-characterized oncogenic drivers, and MET inhibitors could successfully treat several tumor types with MET alterations, including gliomas with PTPRZ1-MET fusion. However, the full diversity and prevalence of MET alterations in gliomas are still lacking to accurately identify a subset of patients likely to benefit from MET inhibitor treatment. Here, we interrogated genomic profiles of 1,351 gliomas, and further identify 60 cases harboring MET alterations, including MET fusions and various MET exon skipping events. MET RNA alterations, but not MET amplification, are highly enriched in the secondary glioblastomas (sGBM) with significantly worse prognosis. Further molecular analysis has shown that MET RNA alterations acting an additive effects of MET overexpression are induced in the course of glioma evolution. In vitro and clinical studies indicate cells and patients harboring MET alterations have better response to MET inhibitors. Collectively, these data suggest that a subgroup of gliomas harboring MET alterations likely to have benefit from MET-targeted therapy.

---

Glioma is the most common and lethal, highly heterogeneous type of tumor in the adult brain, and new targets for individualized or molecularly targeted strategies are urgently needed [1–3]. In addition to MET amplification, a variety of MET RNA alterations have also been reported in Pan-cancer research (**Fig. 1a, Supplementary Table 1**). To identify gliomas likely to potential benefit from MET target therapy, we also screened all potential MET alterations in 1,148 newly collected gliomas and 31 glioma patient-derived cells (PDC) from obtained from the Chinese Glioma Genome Atlas Network (CGGA), and 203 gliomas collected from previous published data (**Supplementary Fig. 1a, Supplementary Table 2**). We found a various of MET RNA alterations, including MET fusion genes (34/1,104 cases), MET exon 10 skipping (METex10) (27/1,104 cases), MET exon 14 skipping (METex14) (23/1,104 cases), and MET exon 19 skipping (METex19) (19/1,104 cases) in glioma and patient-derived cells (PDCs; 7/31 cell lines) (**Fig. 1b**). Of them, MET fusion genes included CAPZA2-MET, KCND2-MET, LHFPL3-MET, RSBN1L-MET, ST7-MET, ZM, ZMC7orf55-LUC7L2-MET, ZMELAVL3-MET, ZMIZ2-MET, ZMST7-MET, and ZNF277-MET (**Supplementary Table 2**); Intriguingly, two novel breakpoints of MET identified to fuse to two different breakpoints of other genes in more than one samples (LHFPL3-MET and RSBN1L-MET). METex10 has been previously identified in lung cancer cell lines [4], but firstly identified in glioma (**Supplementary Fig. 1b**); and METex19 was a novel MET alteration (**Supplementary Fig. 1c**). Furthermore, we validated the presence of METex10 and METex19 in clinical glioma samples via gel-based PCR and Sanger sequencing (**Fig. 1c-f, Supplementary Fig. 1d**).

**Figure 1.**
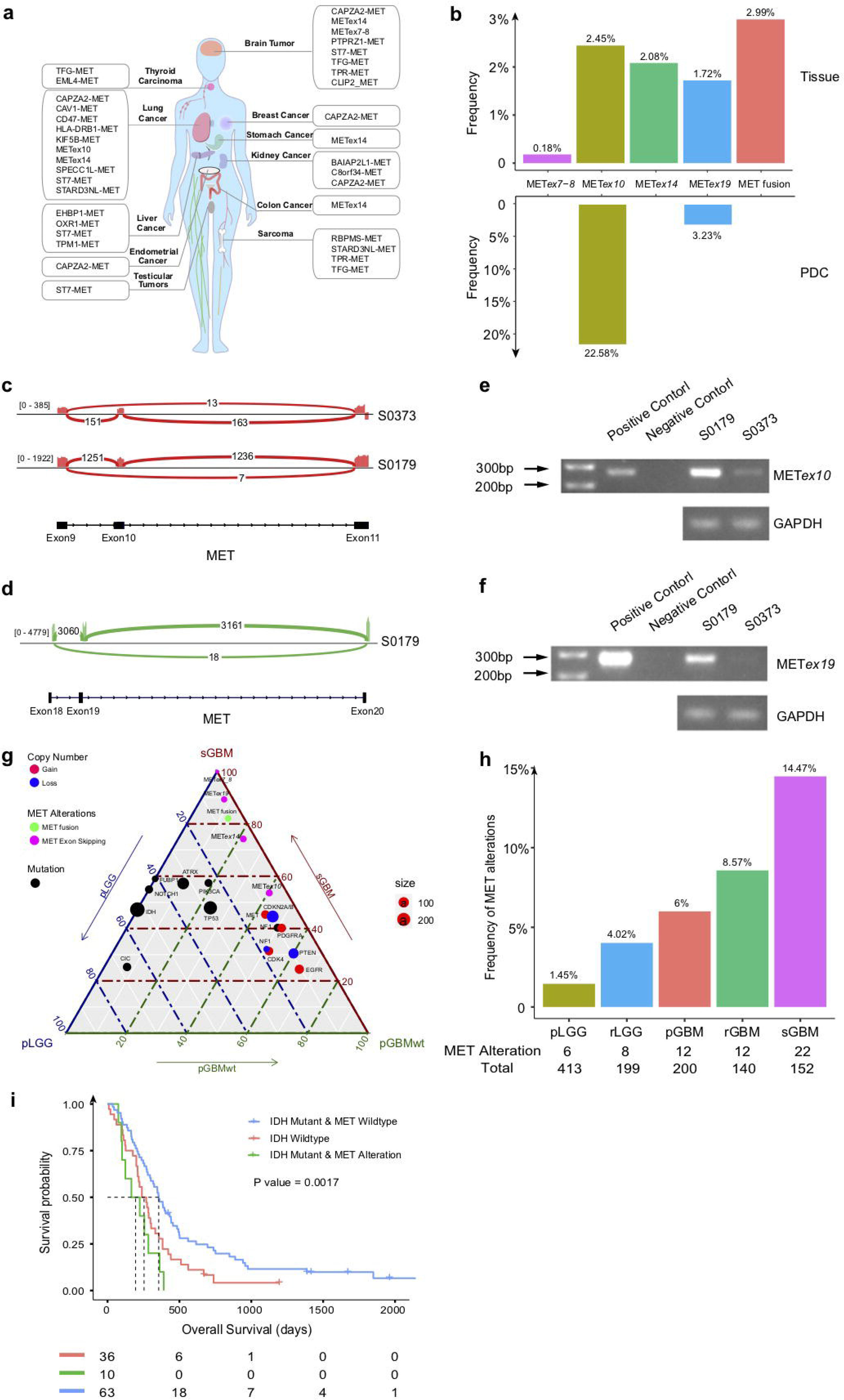
MET alterations frequently occurred and enriched in sGBMs. **(A)** A summary of all reported MET RNA alterations in pan-cancer. **(B)** Frequencies of each MET RNA alterations in glioma subtypes and patient derived glioma cells in this study. **(C-D)** Sashimi plot of RNA-seq data from two METex10-positive sGBM (S0373 and CGGA_S0179, **C**) and a METex19-positive sGBM (S0179, **D**). **(E-F)** Validation of METex10 and METex19 in clinical samples. **(G)** Ternary plot of mutation frequency in driver genes in primary LGG (pLGG), secondary GBM (sGBM), and IDH-wildtype primary GBM (pGBM), and the node size represents their overall frequency in glioma. **(H)** Frequencies of MET alterations in different subtypes of gliomas. **(I)** Overall survival of sGBM patients (first-diagnosis) with and without IDH mutation and MET status (Median survival, 356 days, 254 days, 195 days for IDH Mutation & MET Wildtype, IDH Wildtype, IDH Mutation & MET Alteration, respectively).

We next asked whether METex10 and METex19 play a role in activating MET signaling. Unlike METex14, which induce MET signaling activation though skipping the JM domain of MET protein [8], METex10 and METex19 may have no biological function by themselves because they give rise to truncated MET protein (**Supplementary Fig. 2a,b**). However, the four types of MET RNA alterations, including MET fusion, METex10, METex14, and METex19, were highly enriched in secondary GBM (sGBM) (**Supplementary Fig. 2c–f**). Similarly, comparison of genomic alterations in primary LGG (pLGG), IDH-wild-type primary GBM (pGBM), and sGBM revealed a high frequency of these MET RNA alterations in sGBM (**Fig. 1g**). Interestingly, MET amplification and CDKN2A/2B homogenous deletion were equally distributed in pGBM and sGBM, suggesting that MET RNA alterations may differed from the MET amplification in DNA.

We next compared the mutational landscape of pLGG and sGBM to identify the molecular features of gliomas with different MET alterations. We noticed that different MET RNA alterations could simultaneously present in the same case, and often co-occurred with CDKN2A/2B homogenous deletions (Fisher test, p=2.924e-16) in sGBM (**Supplementary Fig. 2g**). The CDKN2A/2B homogenous deletion is a prognostic marker in patients with IDH-mutant astrocytic gliomas and other types of cancer, Gliomas with lower grade histological features could be define as high grade when with CDKN2A/2B homogenous deletions[5–9]. However, few inhibitors are available for CDKN2A/B-deficient tumors. It has been indicated that RNA interference targeting of MET in CDKN2A-deficient tumor could significantly delays tumor growth [10]. Since MET alterations were correlated with CDKN2A/B homozygous deletion in sGBM, MET maybe comprise a new therapeutic target for patients with CDKN2A/B homozygous deletion.

We next asked whether all different MET RNA alterations were linked to sGBM patient survival. We defined the sGBM overall survival (OS) as the period from the first diagnosis of sGBM until patient death. Similar to METex14 and MET fusion, METex10 and METex19 separately stratified the OS of these patients (**Supplementary Fig. 2h–k**). This suggested that METex10 and METex19 might be “quiescent biomarkers” [11], i.e., marking the same subgroup of cases as METex14 and MET fusion.

To further explore the clinical implications of these MET RNA alterations gliomas, we found that MET RNA alteration is highly enriched in sGBM (**Fig. 1h**). We also observed that the OS of patients with IDH-mutant sGBM with MET alterations was significantly poorer than that in patients with sGBM without MET alterations and with IDH wild type (**Fig. 1i**). This meant that MET RNA alterations could serve as a biomarker to stratify the survival of sGBM.

Given that sGBM indicates tumor progression to a late stage, we inferred the high enrichment of MET alterations in sGBM to suggest that MET alterations might be associated with the secondary activation of MET at a later stage of glioma evolution [1, 3, 12]. Recently, we have reported a novel bioinformatics tools for efficient prediction of the evolution stage of glioma samples based on their transcriptional features. We used this tool to calculate the evolution stage of tumors with different molecular features in **295** glioma samples for which both genetic and transcriptomic data were available. Indeed, gliomas with MET alterations were at a late stage of tumor evolution (Fig. 2a), consistent with the observation that MET alterations were enriched in sGBM (**Supplementary Fig. 3a**).

**Figure 2.**
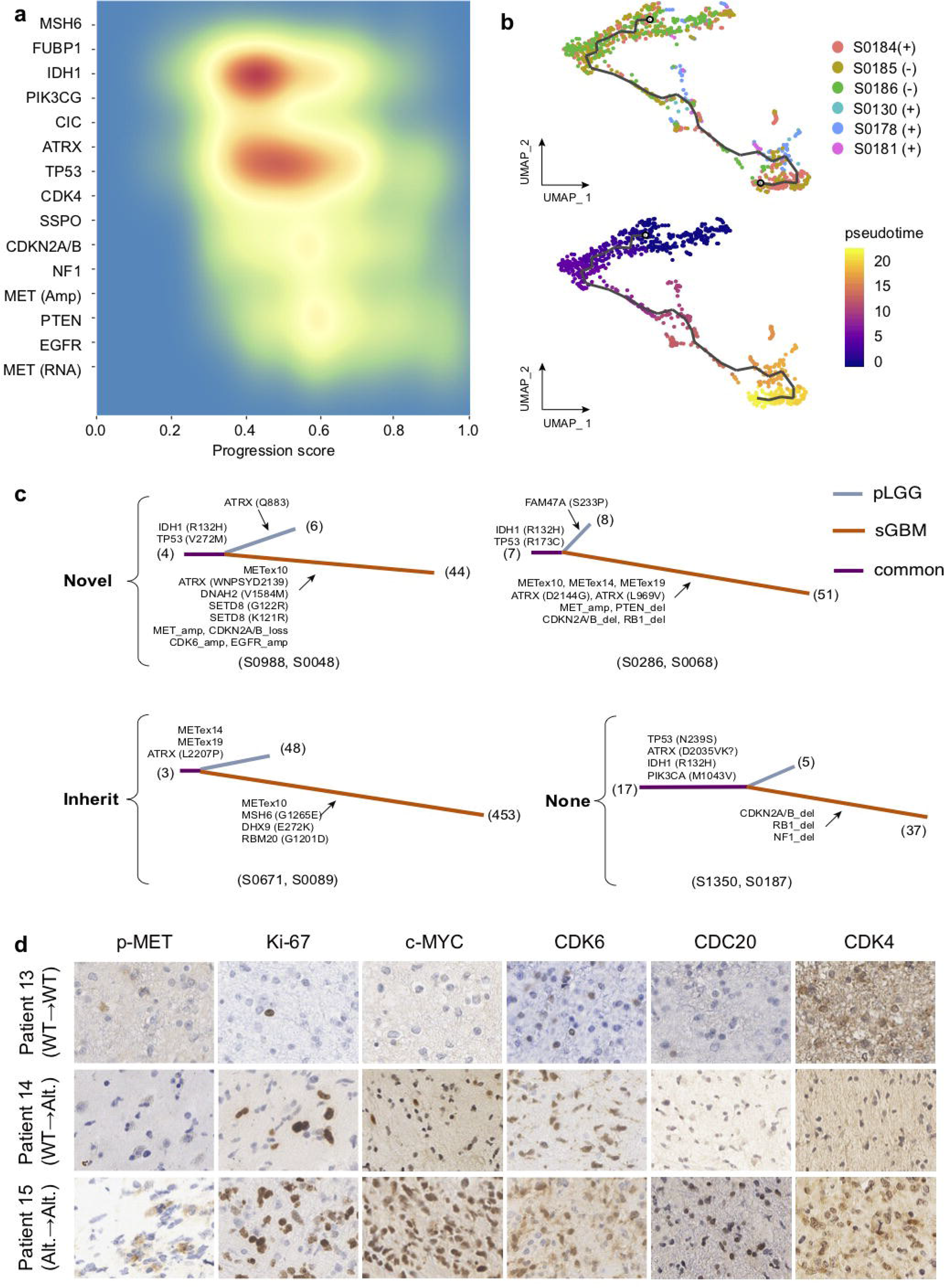
MET alterations are later events. **(A)** Distribution of progression scores of key molecular events in gliomas (n=291). (**B**) The single-cell trajectory analysis (top: UMAP cluster; bottom: pseudotime). **(C)** Evolutionary trees of four patients (two with novel MET alterations occurred in the recurrent samples, one with inherit MET alterations from pre-recurrent samples, and one without MET alterations in both the initial and recurrent samples) evaluated by whole-exome or whole-genome sequencing. Selected driver mutations are labeled in black. **(D)** IHC of p-MET, Ki-67, c-MYC, CDK6, CDC20 and CDK4 from primary glioma with different MET status. Bar=50 μm.

We next performed single-cell RNA-seq (scRNA-seq) analysis of six sGBM samples to study the potential evolution trajectory of tumors with MET RNA alteration (**Supplementary Fig. 3b**). Using a single-cell trajectory analysis, malignant cells in four cases with ZM were in the late stage of pseudotime cancer development (**Fig. 2b**). Collectively, these findings indicated that MET alterations evolve with tumor progression.

It has been reported that IDH-mutant non-co-deletion cases with CDKN2A/B homozygous deletion are associated with genomic instability [2, 13]. Here, we also noticed that, IDH1; TP53, ATRX, CDKN2A/B homozygous deletion, RB1 alterations were also significantly co-occurrence with MET RNA alterations in sGBM (**Supplementary Fig. 3c**). In addition, analysis of longitudinal whole-exon or whole-genome sequencing data revealed that the occurrence of MET alterations was accompanied with the appearance of mutations in DNA damage repair genes in three paired cases with MET RNA alterations, but not in paired samples from a case without MET RNA alterations (**Fig. 2c**). Collectively, this suggests that abnormal DNA damage repair is associated with the occurrence of MET alterations during tumor evolution.

It remains an open question whether distinct clinical outcomes can be observed depending on the evolutional patterns, to predict tumor progression [14]. Here, we analyzed longitudinal RNA-seq data of 8 paired samples to explore features associated with occurrence of MET alterations in the secondary GBM. We observed that the read number ratios which supporting MET RNA alteration is dramatically elevated during glioma progression in all eight patients (n=6, progression from LGG to sGBM; and n=2, progression from sGBM to sGBM) (**Supplementary Fig. 3d**). Importantly, MET alteration is also present in sGBM-paired initial tumor diagnosed as LGG (WHO grade II-III), indicating MET RNA alteration in LGG could be used to predict the occurrence of MET RNA alteration in the sGBM.

To further dissect which features of LGG are associated with their evolution into sGBM with MET RNA alteration, we studied the transcriptional characteristics of all LGG with MET alterations. We observed that both, MET mRNA levels and progression scores of LGG harboring MET alterations exceeded the respective median values of all LGG patients (**Supplementary Fig. 4a,b**). GSEA analysis confirmed enrichment of multiple hallmark pathways associated with cell-cycle progression, DNA replication, and transcription in LGG samples with MET RNA alterations (**Supplementary Fig. 4c**).

Next, we also compared the transcriptional features of LGG that ultimately evolved to sGBM harboring MET alterations with those that evolved to sGBM with wild-type MET. We found that the initial gliomas that tended to evolve to sGBM harboring MET alterations were enriched for hallmarks of mitotic spindle, G2M checkpoints, and E2F and MYC target genes (**Supplementary Fig. 4d**). This is consistent with the notion of a replication-stress phenotype in tumors [15]. We validated these findings by IHC (**Fig. 2d**). Specifically, the initial glioma that finally evolved to sGBM with MET alterations exhibited higher expression of c-MET, p-MET, Ki-67, CDK6, CDK4, DNA instability marker CDC20 [16], and MYC than the initial glioma that finally evolved to MET wild-type sGBM, and we defined this feature as IHC characteristic of the initial tumors that tended to evolve to sGBM. Collectively, these observations indicate that LGGs with high DNA replication stress status, defined by enhanced cell proliferation and transcriptional activities, trend to evolve to sGBM with MET RNA alterations.

MET overexpression can activate cell proliferation, reduce apoptosis, and promote tumor migration[17, 18]. Consequently, MET overexpression is expected to be strongly associated with MET alterations. However, MET overexpression does not always accompany MET amplification or METex14 in lung cancers [19]. Here, we investigated the association between MET overexpression and the MET alterations in sGBM. We observed that MET mRNA levels were significantly increased in sGBM with MET RNA alteration (**Fig. 3a**). That was also the case in sGBM with each MET RNA alteration variant (**Supplementary Fig. 5a–d**). Interesting, MET expression significantly increased with an increasing number of MET alteration events (**Supplementary Fig. 5e**). Further, as determined by immunohistochemistry with specific antibodies, total MET (c-MET) and phosphorylated MET (p-MET) levels were elevated in sGBM with MET RNA alterations, compared with sGBM without MET RNA alterations (**Fig. 3b**). Collectively, these observations suggest that MET RNA alteration is a robust biomarker to predict the MET overexpression in sGBM.

**Figure 3.**
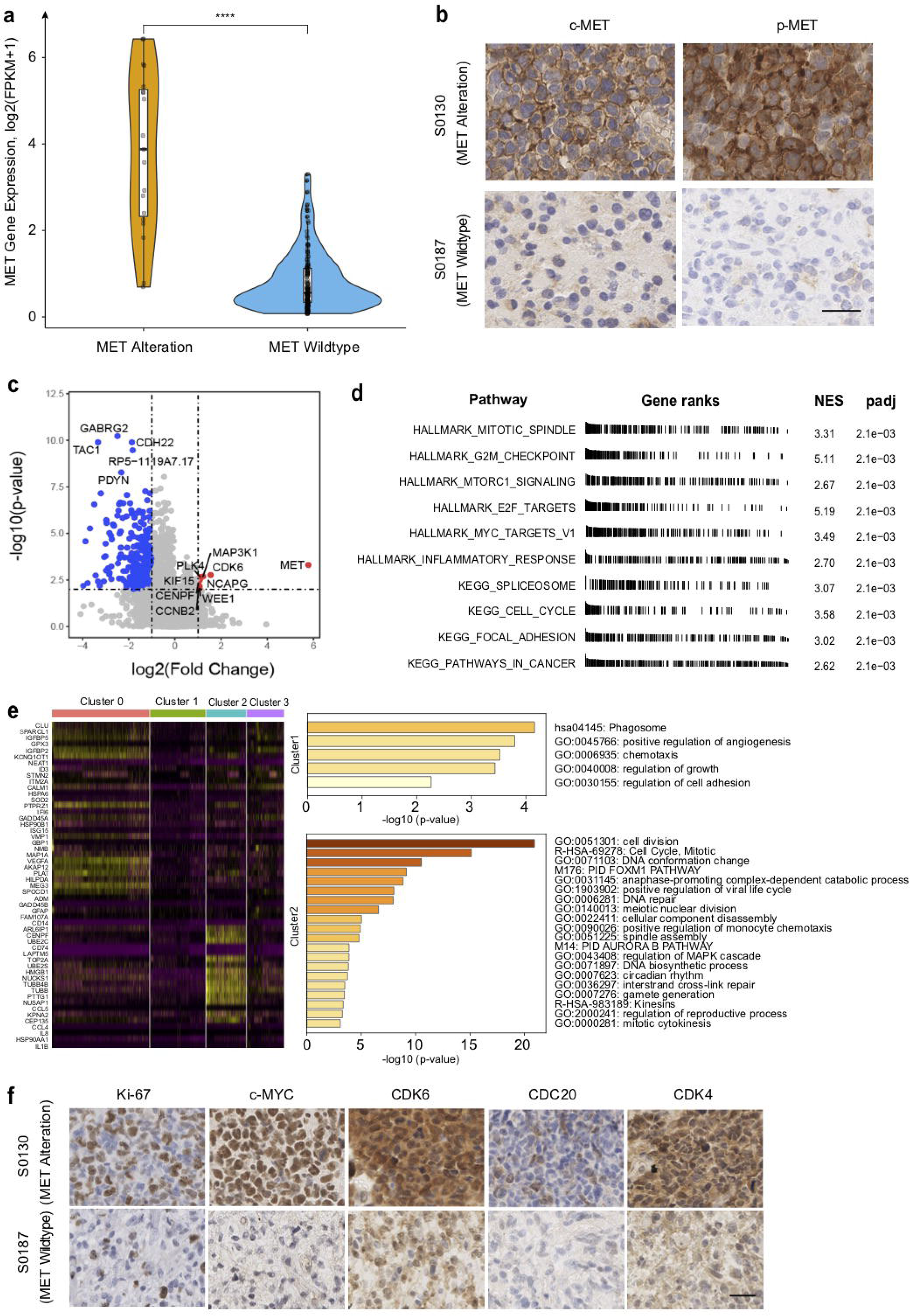
Alternative pathways in sGBM with MET alterations. (**A**) The distribution of MET expression in glioma with MET alterations and MET wildtype. **** P<0.0001 in Wilcoxon Rank Sum Test. (**B**) The IHC of cMET /pMET in sGBM with/without MET alterations. Bar=50 μm. **(C)** Volcano plot showing fold change and the adjusted p value of genes differentially expressed between MET with and without alterations. **(D)** The most enriched cancer hallmarks and KEGG pathways for applying the gene set enrichment analysis (GSEA) in sGBM cases with MET alterations versus that without MET alterations. **(E)** The relative expression level of genes across cells is shown, sorted by cell type. MET alterations are enriched in cluster 2 and cluster3. The GO and KEGG pathway analysis of cluster 2 and cluster3 are shown. **(F)** The IHC of Ki-67, p-MYC, CDK6, CDC20 and CDK4 in sGBM with and without MET alterations. Bar=50 μm.

To further examine whether sGBM harboring MET RNA alteration also presents biological characteristics which could be triggered by MET signaling activation. We compared gene expression profiles in MET wild-type sGBM samples with those in sGBM samples with MET RNA alterations. We identified 672 genes that were up-regulated in the latter, with MET the most significantly up-regulated gene (**Fig. 3c**). GSEA analysis revealed an enrichment of multiple hallmarks and pathways associated with cell-cycle progression, DNA replication, and transcription in these samples (**Fig. 3d**). In addition, as per scRNA-seq analysis, malignant tumor cells were clustered into four clusters (clusters 0–3), and the cluster 1 and 2 cells were significantly more numerous in tumors harboring ZM fusion than in MET wild-type tumors (**Fig. 3e and Supplementary Fig. 5f**). Functional analysis revealed enrichment of cell adhesion and proliferation activities in clusters 1 and 2, respectively (**Fig. 3e**), suggesting that ZM fusion were associated with increased cell proliferation and migration. We validated these transcriptomic findings by IHC (**Fig. 3f**). Finally, we observed that the proliferation rate of PDCs harboring METex10 and METex19 (PDC_25-1) was higher than that of PDCs with MET wild type (PDC_22-1) (**Supplementary Fig. 5g**). Taken together, the above observations suggest that MET alterations could be indicative of the DNA replication stress status, defined by enhanced cell proliferation and transcriptional activities, in sGBM.

To study the characterization of MET alterations on drug response, we investigated drug sensitive test on response to TMZ and a MET inhibitor Crizotinib in PDCs. We observed that PDC_25 and PDC_28 harboring MET alterations exhibited less sensitive to TMZ than PDC_18 and PDC_22 (**Fig. 4a**), consistent with our previous report that gliomas with high MET expression are resistant to TMZ [20]. On the other hand, PDC_25 and PDC_28 were more sensitivity to crizotinib, an FDA-approved MET-targeted drug, compared with MET wild-type PDCs (PDC_18 and PDC_22) (**Fig. 4b**). The findings indicated that gliomas with MET alterations might be sensitive to MET-targeted therapy.

**Figure 4.**
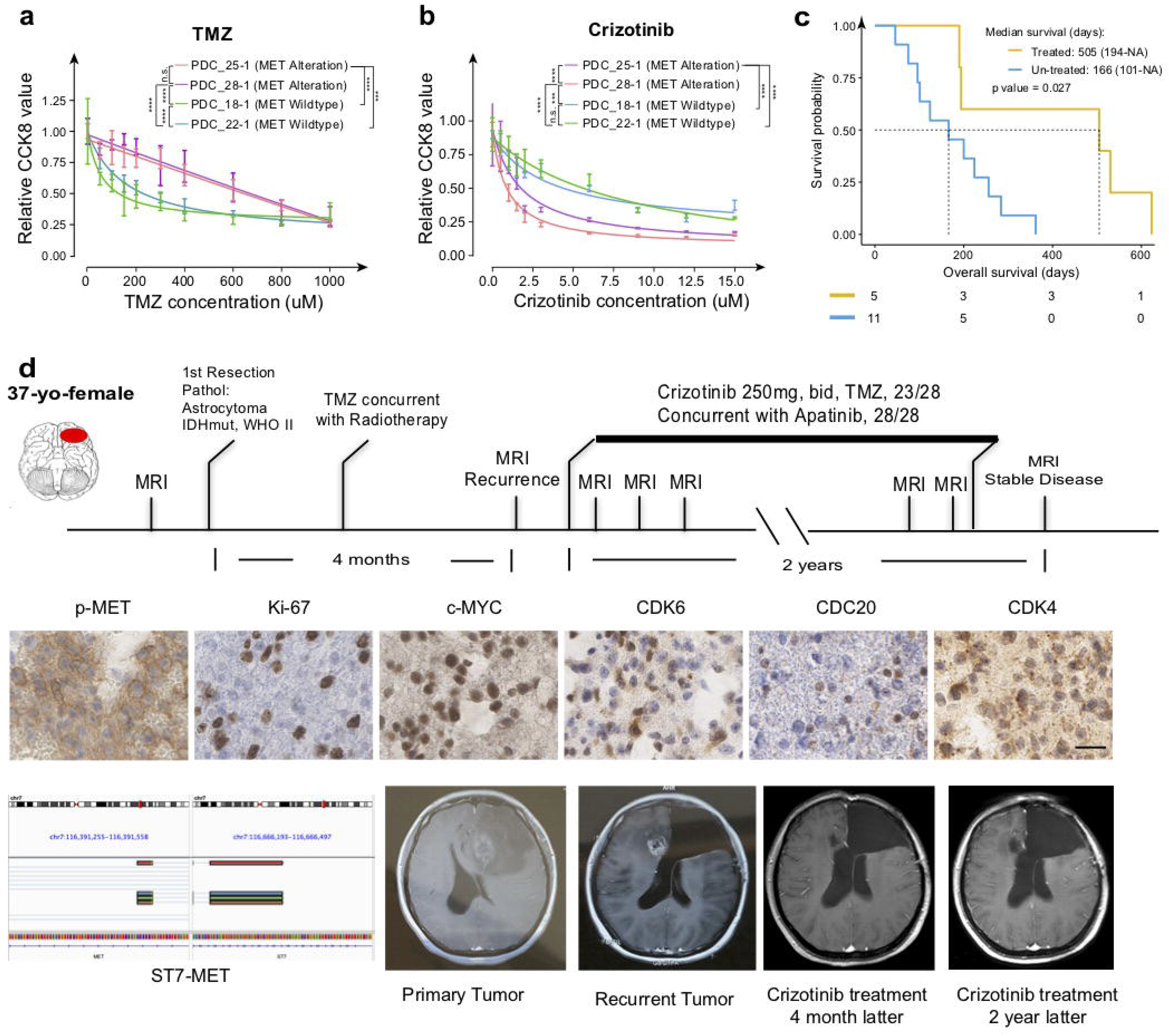
Preclinical testing and translation of a MET inhibitor treatment into a clinical setting. **(A-B)** The relative viability of cells (represented by the relative CCK8 value) after treatment of crizotinib (**A**) and TMZ (**B**). **(C)** Overall survival of sGBM patients (with MET alterations) with or without MET inhibitors (PLB1001). **(D)** Diagram of the glioma progression and treatment in patient S1451. IHC of p-MET, Ki-67, c-MYC, CDK6, CDC20 and CDK4 from primary glioma sample of this patient. Bar=50 μm. All MRI images represent axial T1-weighted scans with contrast enhancement of the patient at baseline and at indicated time points after initiation of treatment. Crizotinib (250 mg bid) concurrent with apatinib (850mg qd) were administrated orally at recurrence after 4 months from the diagnosis. Marked tumor shrinkage at the site of the main lesion was observed (arrows).

In our clinical trial (NCT02978261) studying of a c-Met inhibitor PLB1001 in glioma patients, all of five sGBM patients harboring MET RNA alterations who had received MET-specific small-molecule inhibitor PLB-1001 have reached the end of follow-up. The OS of these five patients with MET inhibitor treatment significantly improved, compared with OS of other eleven sGBM patients with MET RNA alterations without MET inhibitor treatment in our cohort (**Fig. 4c**), suggesting patients with MET alterations conferred more sensitivity to MET-targeted therapy.

Encouragingly, we found that a recurrent glioma patient whose primary glioma with MET RNA alteration (ST7-MET) well response to MET-targeted therapy after resistant to standard chemoradiotherapy. This patient (S1351) was a 37-year-old female, diagnosed with astrocytoma with IDH mutation, WHO grade II, which was resected followed by TMZ concurrent with radiotherapy. A MET fusion was detected in the specimen by RNA-sequencing. The initial tumor also showed a IHC characteristics of the initial tumors that tended to evolve to sGBM with MET alterations (**Fig. 4d**). The patient later underwent crizotinib treatment (250 mg bid) concurrent with apatinib (850 mg qd) at recurrence 4 months from the diagnosis. Tumor shrinkage and symptom relief were observed after 4 months of MET inhibitor treatment. The patient is still alive, with an over 2-year post-progression survival (**Fig. 4e**). This indicates that the MET inhibitor treatment is beneficial to not only sGBM patients who harbor ZM and/or METex14, but also to those with other MET RNA alterations (ST7-MET). Considering the similarity among various MET RNA alterations identified in this study, MET inhibitor, as recently described in many types of cancer, might represent a promising therapeutic option for glioma patients with these MET RNA alterations.

In conclusion, we have highlighted the characteristics of MET RNA alterations in glioma. We also demonstrated that MET RNA alterations were induced in the course of glioma evolution. In addition, MET RNA alterations were additive effects of MET overexpression and were strongly associated with aggressive biological phenotypes in glioma. Clinically, these RNA alterations could serve as biomarkers to predict whether glioma would benefit from MET-targeted therapy.

## Methods

### Sample acquisition

Glioma tissues, the corresponding genomic data, and patients’ follow-up information (histology, WHO grade, gender, age, extent of resection, and OS) were obtained from the CGGA, which includes patient samples collected from Beijing Tiantan Hospital, Sanbo Hospital in Beijing, Tianjin Medical University General Hospital, The First Affiliated Hospital of Nanjing Medical University, Harbin Medical University, and China Medical University. The glioma tissues were snap-frozen in liquid nitrogen immediately after surgical resection and preserved in liquid nitrogen.

### Sample processing and sequencing

Genomic DNA from tumors and matched blood samples was extracted and confirmed DNA integrity by 1% agarose gel electrophoresis. The DNA was subsequently fragmented, and underwent quality control. Then, pair-end libraries were prepared. For whole-exome sequencing, Agilent SureSelect kit v6 was used for target capture. The sequencing was done using Illumina Hiseq platform with a 150-bp pair-end sequencing strategy. Total RNA from each tumor sample was extracted, and then tRNA and rRNA removed. After quality control and quantification, mRNA was reverse-transcribed to cDNA. Next, library was prepared and the samples sequenced.

### Mapping and mutation calling

Valid DNA sequencing data were mapped to the reference human genome (UCSC hg19) using the Burrows-Wheeler Aligner (BWA v0.7.12, bwa mem) with default parameters [21]. Then, SAMtools v1.2 [22] and Picard v2.0.1 (Broad Institute) were used to sort the reads by coordinates and mark duplicates. Statistics such as sequencing depth and coverage were calculated using the resultant BAM files. SAVI2 was used to identify somatic mutations (including single-nucleotide variations and short insertion/deletions), as previously described [23, 24]. In this pipeline, SAMtools mpileup and bcftools v0.1.19 were used for variant calling. Then, the preliminary variant list was filtered to remove positions with no sufficient sequencing depth, positions with only low-quality reads, and positions that were biased toward either strand. Somatic mutations were identified and evaluated by using an Empirical Bayesian method. In particular, mutations with the mutation allele frequency in tumors significantly higher (fisher p value <0.05) than that in normal control were selected.

### Copy number alteration analysis

For whole-exome sequencing data, CNVkit v0.9.4 [25] was used to detect copy number changes.

### Detection of gene fusions in RNA-seq data

STAR-Fusion v1.2.0 (https://www.biorxiv.org/content/10.1101/120295v1) was used to detect gene fusion in RNA-seq data, with default parameters.

### Definition of MET alteration

In this study, we have defined the MET alteration as one (or more) MET RNA alteration (MET fusion, METex10, METex14, and METex19).

### Detection of MET exon skipping in RNA-seq data

In the study, MET 10, 14, and 19 exons skipping were detected based on reads spanning the junction of exons 9 and 11, exons 13 and exon 15, and exon s18 and exon 20, respectively. Briefly, RNA-seq reads were aligned with the reference genome (hg19) using STAR. Then, the reads containing a jump of 4,429 bp from chr7:116398664, 3,226 bp from chr7:116411700, and 1,3666 bp from chr7:116422042 were counted to identify the candidate MET exon 10, 14, and 19 skipping events, respectively. The spanning reads were manually checked in Integrative Genomic Viewer (IGV) to remove false-positives, such as PCR artifacts and potential mapping errors. IGV was used to visualize the skipped reads.

### Validation of MET exon skipping by PCR and Sanger sequencing

RNA extraction and reverse-transcription of the glioma specimens were performed as mentioned. MET exon skipping was detected by PCR. The following primers were used to detect exon 14 skipping: forward: 5’-AATCTTTTATTAGTGGTGGGAGCACAAT-3’; and reverse: 5’-GAATTAGGAAACTGATCTTTAATTTGC-3’. The reverse primer crosses exons 13 and 15 of MET. The following primers were used to detect MET exon 10 skipping: forward: 5’- CAAATCTTTTATTAGGCATGTCAACATC-3’; reverse: 5’-CTTCTGGAAAAGTAGCTCGGTAGTCT-3’. The forward primer crosses exons 9 and 11 of MET. The following primers were used to detect MET exon 19 skipping: forward: 5’-GATCAGTTTCCTAATTCATCTCAGAAC-3’; reverse: 5’- GCCAAAGGACCAATACAGTTTCTT-3’. The reverse primer crosses exons 18 and 20 of MET. In all reactions, DNA polymerase (GoTaq, Promega) was used to amplify a 626-bp fragment, with 30 cycles of 94°C, 30 s; 58°C, 30 s; 72°C, 60s; and a final extension at 72°C for 10 min. The product bands were extracted from agarose gel (1%) after electrophoresis and verified by Sanger sequencing.

### Gene expression analysis and GSEA

The human reference genome hg19 and genome annotation file were downloaded from the UCSC genome browser. Clean RNA sequencing reads were mapped to the reference genome using STAR. Gene expression, in FPKM, was calculated by RESM. Differentially expressed genes were selected based on fold change and adjusted p-value, which was calculated by using DESeq2 [26]. R package fgsea was used for GSEA analysis. Normalized enrichment scores (NES) were calculated for gene sets that were downloaded from MSigDB database version 6.1 (http://software.broadinstitute.org/gsea/msigdb).

### Sample and library preparation for scRNA-seq

Samples were collected from Beijing Tiantan Hospital, Capital Medical University, from six secondary glioblastoma patients. Fresh clinical samples were first trypsinized, followed by isolation of single cell and dead cell removal to obtain cell suspensions. Cell viability reached 80% with cell count reached 5×10^5^ cells. cDNA libraries were prepared using the Chromium Single Cell 3’ Library and Gel Bead kit v2 (120267, 10× Genomics) according to the manufacturer’s instructions. RNA-seq was performed using the Illumina HiSeq 2500 platform.

### Single-cell data processing, quality control, and analysis

The Cell Ranger guideline (v3.0.2, 10× Genomics) was followed to convert Illumina base-call files to FASTQ files, aligning FASTQ files twith the hg19 reference (v3.0.2, 10× Genomics) for human samples, and producing a digital gene-cell counts matrix. Cells in gene-cell matrices with fewer than 200 transcripts, and genes with fewer than two counts in two cells were removed. The matrix was then normalized such that the number of UMIs in each cell was equal to the median UMI count across the dataset and log-transformed. The SMART-seq2 and 10X Genomics datasets were integrated using the IntegrateData function in R package “Seurat V3.0”. To pare-down the gene expression matrix to the most relevant features, principal components analysis (PCA) was used; this changes the dimensionality of the dataset. FindClusters function was performed to generate four clusters. To visualize data in 2D space, the PCA-reduced data were analyzed by Uniform Manifold Approximation and Projection (UMAP), a non-linear dimensional reduction method. The top markers for each cluster were plotted using the DoHeatmap function. The differentially expressed genes between clusters were screened using the FindMarkers function and a comprehensive biological functional annotation of the gene list was performed in Metascape (https://metascape.org/). Cell subtypes (malignant and benign cells) were identified by cell-specific markers. All malignant cells were ordered in pseudotime and single-cell trajectories were built using the R package “Monocle3”.

### Cell culture

Patient-derived cells (PDC) were obtained from fresh surgical specimens of human primary GBM and cultured as tumor spheres in DMEM/F12 medium supplemented with B27 supplement (Life Technologies), bFGF and EGF (20 ng/ml each), at 37 °C in a humidified atmosphere of 5% CO_2_.

### Determination of cell sensitivity to drugs

Cell drug sensitivity was evaluated by using the CCK-8 kit (Dojindo Laboratories, Kumamoto, Japan) according to the manufacturer’s protocol. Briefly, 3,000 PDCs were grown in each well of 96-well plate in normal medium for 24 h. The cells were then incubated in a medium containing specific concentrations of TMZ or crizotinib for 72 h, with six repeats for each concentration tested. Next, 10 μl of CCK-8 reagent was added to each well, and sample optical density at 450 nm and 630 nm was determined after 2 h incubation and 37°C in a humidified atmosphere of 5% CO_2_. The number of living cells was calculated as OD450–OD630.

### Immunohistochemistry

Immunohistochemical staining was performed as previously described [27, 28]. Briefly, tissue sections were deparaffinized and boiled in citrate antigen-retrieval buffer. Then, the primary antibodies (anti–c-MET, 1:1000, Abcam, Cambridge, UK; anti–p-MET, 1:200, CST, MA, USA; anti–Ki-67 1:100, ZSGB-Bio, anti-CDK4, 1:150 dilutions, Proteintech, China; anti-MYC, 1:150, Abcam; anti-CDK6, 1:150, Abcam; anti-CDC20,161 1:150, Abcam) were incubated with the sections, overnight at 4°C. After incubation with ethanol containing 3% hydrogen peroxide for 10 min and washing with PBS buffer, the sections were incubated with the secondary antibodies (1:100, ZSGB-Bio, Beijing, China) at 25°C for 1 h. The images were captured by using Axio Imager 2 (Zeiss) after 3,3’-diaminobenzidine and hematoxylin staining.

### Statistical analysis

Statistical analyses were performed by unpaired two-tailed Student’s T-test or Wilcoxon rank-sum test. Surival curves were estimated with the Kaplan-Meier method. All statistical analyses. Were conducted and obtained using the R software (https://www/r-project.org).

## Acknowledgements

This project was supported by grants from the Natural Science Foundation of China (NSFC)/Research Grants Council (RGC), Hong Kong, China Joint Research Scheme (81761168038); the National Natural Science Foundation of China (81802994, 81972816).

## Author Contributions

J.T., Z.Z. and Q.W. conceptualized the project. Z.Z., Z.B., R.C. and L.W. analyzed and interpreted the sequencing data. J.C., R.C., K.Z., and H.L. performed the experiments. G.L., Y.C. and Q.W. prepared patient samples for sequencing and partially performed data pre-processing. R.T. carried out the clinical patients with MET inhibitor. Q.M., H.H., F.Z., Z.W., F.W., Y.Z. and Y.L. undertook the prospective patient enrolment, diagnosis, and clinical follow-up. J.W., W.Z. and C.K. and guided data analysis. Z.B., R.C. and Z.Z. wrote the manuscript, which was further revised by T.J. and Q.W. All authors have read and approved the manuscript.

---

**Figure S1.**
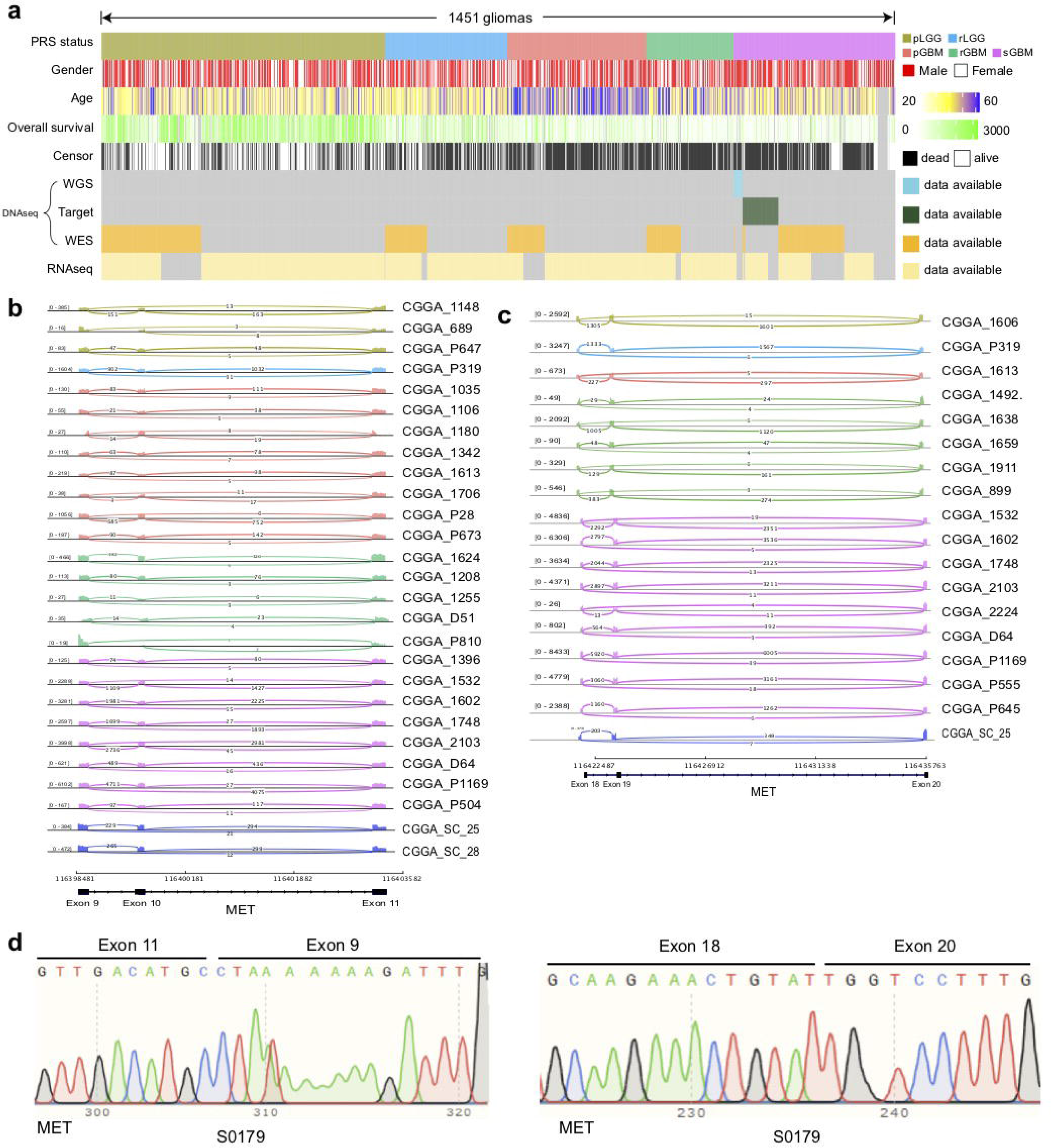
Data summary and validation of MET alterations. **(A)** The data composition of this study. **(B-C)** Sashimi plot of RNA-seq data of all cases with METex10 and METex19 (**C**). **(D)** Sanger sequencing of PCR products of case S0179 with METex10 and METex19.

**Figure S2.**
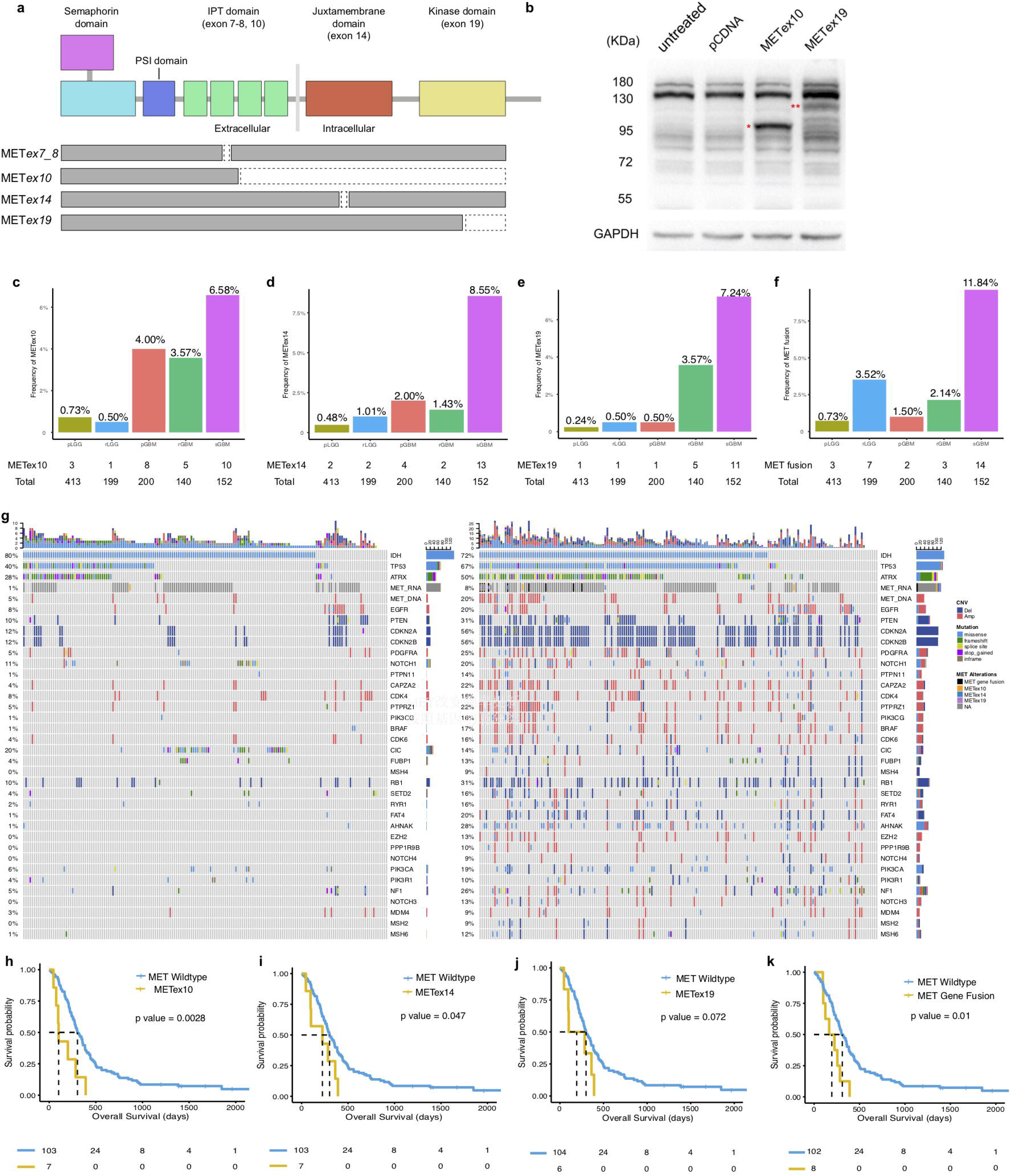
Characteristic of METex10 and METex19. **(A)** Diagram of the protein products of different exon skipped MET. **(B)** The truncated protein band of METex10 (single star) and METex19 (double stars). **(C-F)** Frequencies of METex10, METex14, METex19, and MET fusions in pLGG, rLGG, pGBM, rGBM and sGBM. **(G)** The mutational landscape of pLGG and sGBM. **(H-K)** Overall survival of sGBM patients (first-diagnosis) stratified by the METex10 (H, median survival, 101 days and 303 days for METex10 and non-METex10), METex14 (I, median survival, 224 days and 30 days for METex14 and non-METex14), METex19 (J, median survival, 192.5 days and 299.5 days for METex19 and non-METex19), and MET fusions (K, median survival, 195 days and 310.5 days for MET fusion and non-MET fusion).

**Figure S3.**
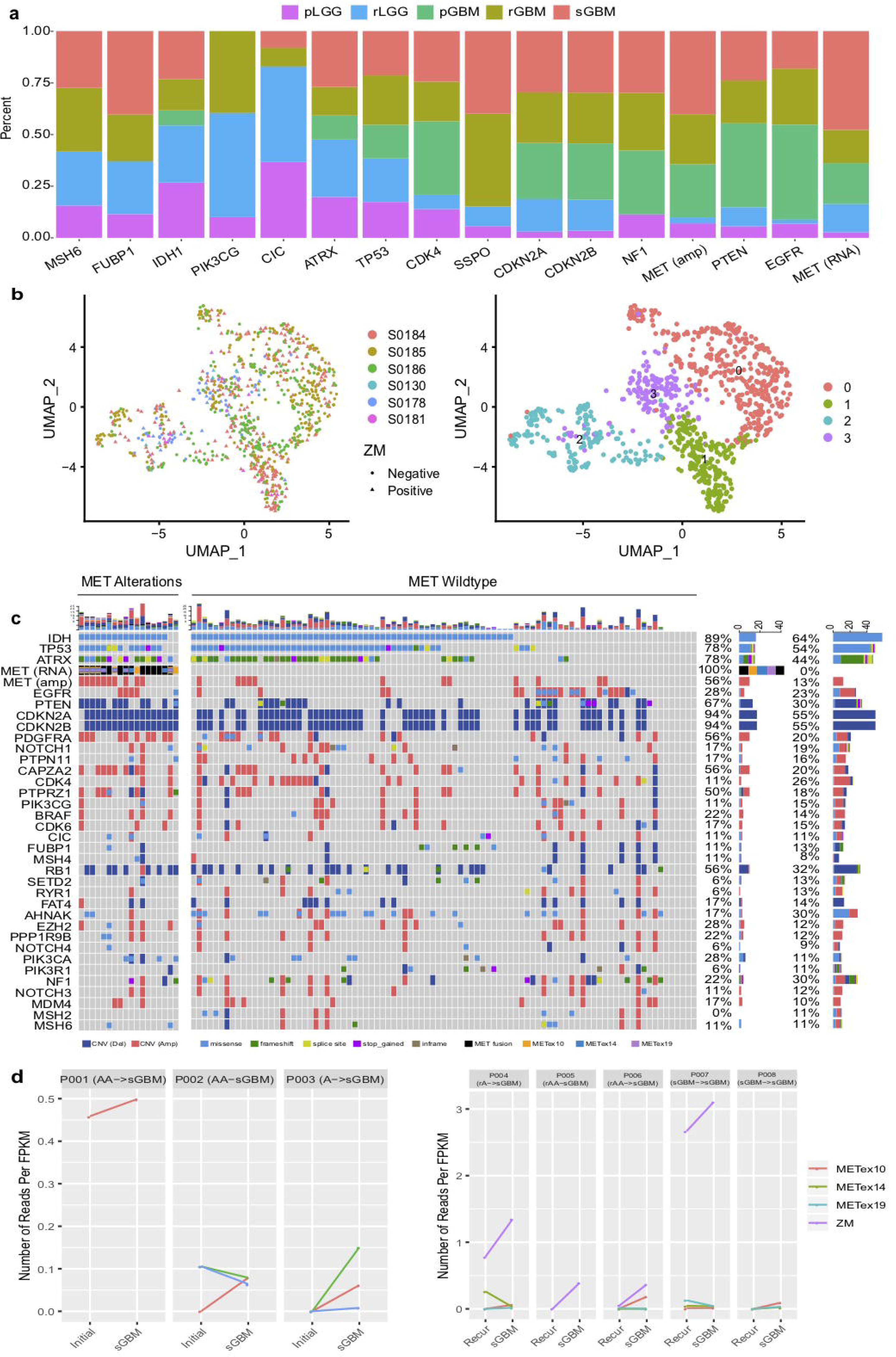
MET alterations are results of tumor evolution. **(A)** Distributions of key molecular events in different gliomas. (**B**) Single cell clusters of sGBM tumors with or without ZM fusion. **(C)** The mutational landscape of sGBM with or without MET alterations. **(D)** Number of MET alteration reads in 8 patients with paired initial tumors and progressed sGBM available. FRPM (Fragments Per Kilobase of Transcript per Million Mapped Reads).

**Figure S4.**
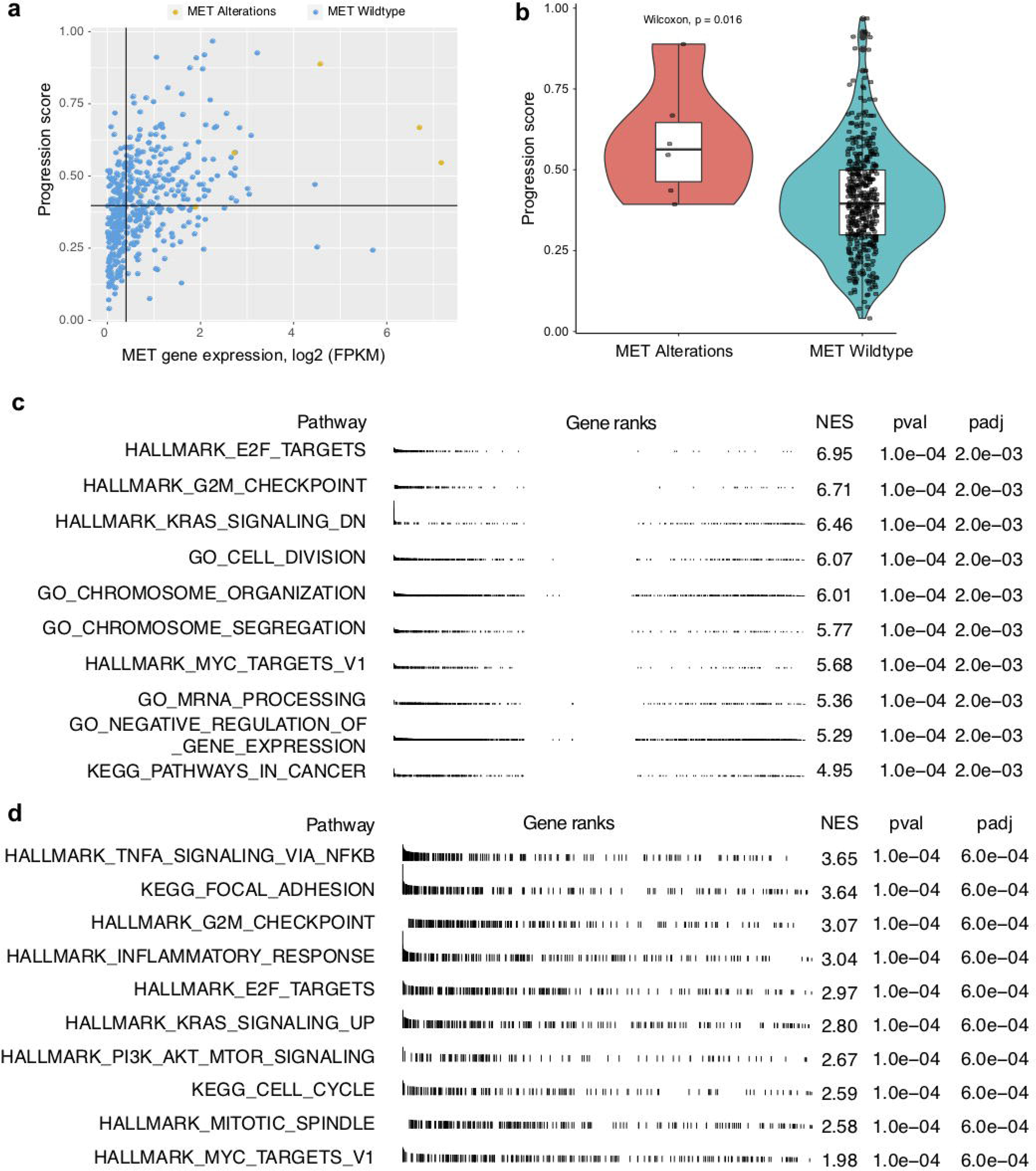
Characteristics of primary gliomas with MET alterations or evolved to sGBM with MET alterations. **(A)** The distribution of MET mRNA expression and progression scores of primary LGGs with or without MET alterations. **(B)** The comparison of progression scores of primary LGGs with or without MET alterations. **(C)** The most enriched cancer hallmarks and KEGG pathways for applying the gene set enrichment analysis (GSEA) in pLGG cases with MET alterations versus that without MET alterations. **(D)** The most enriched cancer hallmarks and KEGG pathways for applying the gene set enrichment analysis (GSEA) in pLGG cases which developed to sGBM with MET alterations versus those pLGG cases which developed to sGBM without MET alterations.

**Figure S5.**
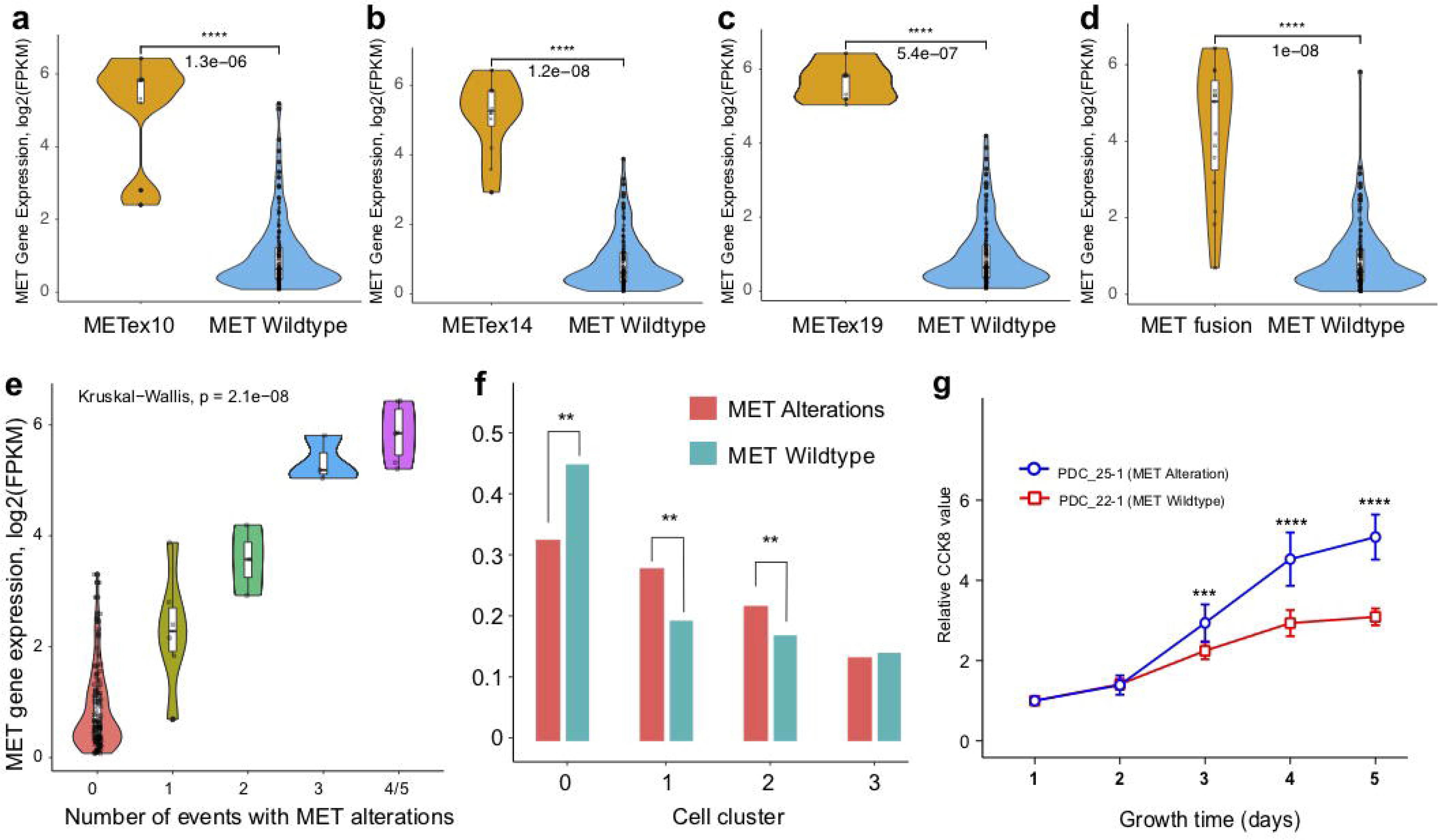
Characteristics of gliomas with MET alterations. (**A-D**) The distribution of MET expression in sGBM with or without METex10 (**A**), METex14 (**B**), METex19 (**C**), and MET fusions (**D**). **(E)** The distribution of MET expression in sGBM with different type of MET RNA alteration events. **(F)** The comparison of frequencies of malignant cells form sGBM with or without ZM fusion in 4 clusters of single-cell transcriptional analysis. **(G)** The proliferation rate of patient derived cells with or without MET alterations.

